# DMSO potentiates the suppressive effect of dronabinol, a cannabinoid, on sleep apnea and REM sleep

**DOI:** 10.1101/769463

**Authors:** Michael W. Calik, David W. Carley

## Abstract

**Purpose:** Dimethyl sulfoxide (DMSO) is an amphipathic molecule with innate biological activity that also is used to dissolve both polar and nonpolar compounds in preclinical and clinical studies. Recent investigations of dronabinol, a cannabinoid, dissolved in DMSO demonstrated decreased sleep apnea frequency and time spent in REM sleep in rats. Here, we tested the effects of dronabinol dissolved in 25% DMSO (diluted in phosphate-buffered saline) to rule out potentiating effects of DMSO.

**Methods:** Sprague-Dawley rats were anesthetized and implanted with bilateral stainless steel screws into the skull for electroencephalogram recording and bilateral wire electrodes into the nuchal muscles for electromyogram recording. Each animal was recorded by polysomnography. The study was a fully nested, repeated measures crossover design, such that each rat was recorded following each of 8 intraperitoneal injections separated by three days: vehicle (25% DMSO/PBS); vehicle and CB_1_ antagonist (AM 251); vehicle and CB_2_ antagonist (AM 630); vehicle and CB_1_/CB_2_ antagonist; dronabinol (CB_1_/CB_2_ agonist); dronabinol and CB_1_ antagonist; dronabinol and CB_2_ antagonist; and dronabinol and CB_1_/CB_2_ antagonist. Sleep was manually scored into NREM and REM stages, and apneas were quantified.

**Results:** Dronabinol dissolved in 25% DMSO did not suppress apnea or modify sleep efficiency compared to vehicle controls, in contrast to previously published results. However, dronabinol did suppress REM sleep, which is in line with previously published results.

**Conclusions:** Dronabinol in 25% DMSO partially potentiated dronabinol’s effects, suggesting a concomitant biological effect of DMSO on breathing during sleep.

## Introduction

Dimethyl sulfoxide (DMSO; (CH_3_)_2_SO) is an amphipathic molecule used to dissolve both polar and nonpolar compounds in preclinical and clinical studies [1,2]. Moreover, DMSO is known to increase the bioavailability of lipophilic drugs [3–5] and is widely distributed throughout the body, including the brain [6,7].

Although most often used as a solvent, existing evidence convincingly demonstrates that DMSO has innate biological activity that may confound experimental results when it is used as a solvent for drug delivery. For example, DMSO is known to decrease the integrity of the blood-brain barrier (BBB) [8], block fast axonal transport in the vagus nerve [9], and modulate morphine-induced antinociception [10]. Further, DMSO induces hypothermia [11], reduces pulmonary ventilation [12], enhances hippocampal-dependent spatial memory accuracy, exerts anxiogenic [13] and antiepileptic [14] effects, and changes sleep architecture[15]. DMSO has also been shown to decrease the occurrence of spontaneous type 1 diabetes by modulating the autoimmune response [16].

Administration of dronabinol, a synthetic cannabinoid type 1 (CB_1_) and cannabinoid type 2 (CB_2_) receptor agonist, dissolved in undiluted DMSO (1ml) decreased apnea index and rapid eye movement (REM) sleep in rats [17,18]. Recent experiments using a model of reflex apnea in anesthetized rats implicates the activation of cannabinoid (CB) receptors on the nodose ganglia in the apnea-suppressive effect [19,20], with little impact deriving from CB receptors located in the brain [21].

Taken together, these findings suggest that dronabinol may act to reduce sleep-related breathing disorder by directly activating CB_1_ and/or CB_2_ receptors within the nodose ganglia of the vagus nerve. However, it is unknown if these effects of dronabinol were partially potentiated by the biologically active solvent, DMSO. Here, we report that apneas were not suppressed, but REM sleep was suppressed, by dronabinol dissolved in 25% DMSO; suggesting that DMSO may potentiate the respiratory effects of cannabinoids in a concentration-dependent fashion.

## Materials and Methods

### Animals

Adult male Sprague-Dawley rats (n = 12; ~275 g) purchased from Harlan Laboratories (Indianapolis, IN, USA) were initially housed in duplicate, maintained on a 12:12 hour light:dark cycle (lights on 8:00 am, lights off 8:00 pm) at 22 ± 0.5 °C, and allowed *ad libitum* access to food and water. After surgery, rats were housed singly to prevent loss of headsets. All animal procedures and protocols were approved by the Institutional Animal Care and Use Committee of the University of Illinois at Chicago.

### Surgical Procedures

Implantation of polygraphic headsets has been described before [17,18]. Rats were anesthetized (ketamine:xylazine 100:10 mg/kg; buprenorphine 0.1 mg/kg), stereotaxically immobilized, and implanted with electroencephalographic (EEG) screw electrodes bilaterally threaded into the frontal and parietal bones. Electromyographic (EMG) wire electrodes were implanted in the dorsal nuchal musculature and tunneled subcutaneously to the skull. EEG and EMG leads were soldered to a miniature plastic connector plug (i.e. headset) and affixed to the skull acrylic dental cement. Scalp wounds were closed with Vetbond Tissue Adhesive. Rats were allowed to recover for 7 days before beginning a week of acclimation to handling and plethysmographic recording chambers.

### Polysomnography and Treatment Protocol

Polysomnography (PSG) procedures have been previously described [18]. Rats underwent nine 6-hour PSG recordings, separated by at least 3 days. All recording sessions began at 10:00 and continued until 16:00. Each rat received an IP injection at 09:45. Rats were immediately placed inside a bias-flow-ventilated (2 l/min) whole-body plethysmograph (PLYUNIR/U, Buxco Electronics, Wilmington, DE, USA). A flexible cable was inserted through a narrow “chimney” into the main plethysmography chamber and attached to the rat’s headset. Rats underwent a week of acclimation to handling and to plethysmographic recording chambers, including being connected to the flexible cable. After acclimation, rats were recorded for 6 hours for one occasion prior to the first experimental session to permit adaptation to the recording system, and to assess the quality of EEG and EMG signals. If signal quality was good, then the rats underwent a repeated measures random order crossover design, such that each rat received each of 8 IP injections exactly one time in random order: vehicle alone (25% DMSO in PBS; 1 ml); dronabinol alone (10.0 mg/kg; Mylan Pharmaceuticals, Morgantown, WV); AM250 alone (5.0 mg/kg, Tocris Bioscience, Bristol, UK); AM630 alone (5.0 mg/kg, Tocris Bioscience); or AM251/630 combination (5.0/5.0 mg/kg); or a combination injection (dronabinol/AM251 or dronabinol/AM630 or dronabinol/AM251/AM630). All drugs were dissolved in 25% DMSO in PBS. Respiratory signals were amplified, band-passed filtered (1 to 10 Hz; CyberAmp 380, Axon Instruments, Sunnyvale, CA), and digitized (250 samples/s; Biologic Sleepscan Premier, Natus, San Carlos, CA). EEG and EMG signals were amplified and band-passed filtered (0.5 to 100 Hz and 10 to 100 Hz, respectively) and digitized (250 samples/s; Bio-logic Sleepscan Premier).

Visual scoring was conducted by a blinded and experienced technician. Sleep stages (wake, NREM, and REM) were scored for every 30-second epoch of the 6-hour recording. Wakefulness was characterized by high-frequency and low-amplitude (beta/alpha waves) EEG with high EMG tone.

NREM sleep was characterized low-frequency and high-amplitude (delta waves) and low EMG tone, while REM sleep was characterized by high-frequency and high-amplitude (theta waves) EEG and an absence of EMG tone. Sleep stage percentages, defined as total time spent in a specific sleep stage (awake, NREM, or REM) divided by total time in the plethysmograph, and sleep efficiency, defined as total time spent in sleep (both NREM and REM) divided by total time spent in the plethysmograph, were also quantified.

Apneas were scored as a cessation of breathing for at least 2 seconds, and were quantified as an apnea index (apneas/hour) and separately stratified for overall sleep and NREM sleep. Apneas were further subdivided into post-sigh (preceded by a breath at least 50% larger than the average of the preceding 5 breaths) and spontaneous apneas (not preceded by an augmented breath), and shown as post-sigh and spontaneous apnea indices, respectively [22,23].

### Statistical Analysis

Data (mean ± SEM) were analyzed using IBM SPSS Statistics 22 (New York, NY) mixed model analysis using treatment (CB agonist, CB antagonist, and CB agonist/antagonist interaction) as a fixed effect and animal as a repeated measure, followed by post hoc multiple comparison tests with Sidak’s correction if there were significant main effects or a significant interaction of main effects. Repeated covariance structure was chosen according to the best-fit Schwarz’s Bayesian information criterion [24].

## Results

Rats (N = 12) were injected with a CB receptor agonist (dronabinol) or vehicle, and with CB_1_/CB_2_ receptor antagonists (AM251, AM630, or both) or vehicle. Sleep efficiency is depicted in Fig. 1. Stratified apnea indexes are presented in Fig. 2, and time spent in wakefulness, NREM or REM sleep is shown in Fig. 3.

**Figure 1.**
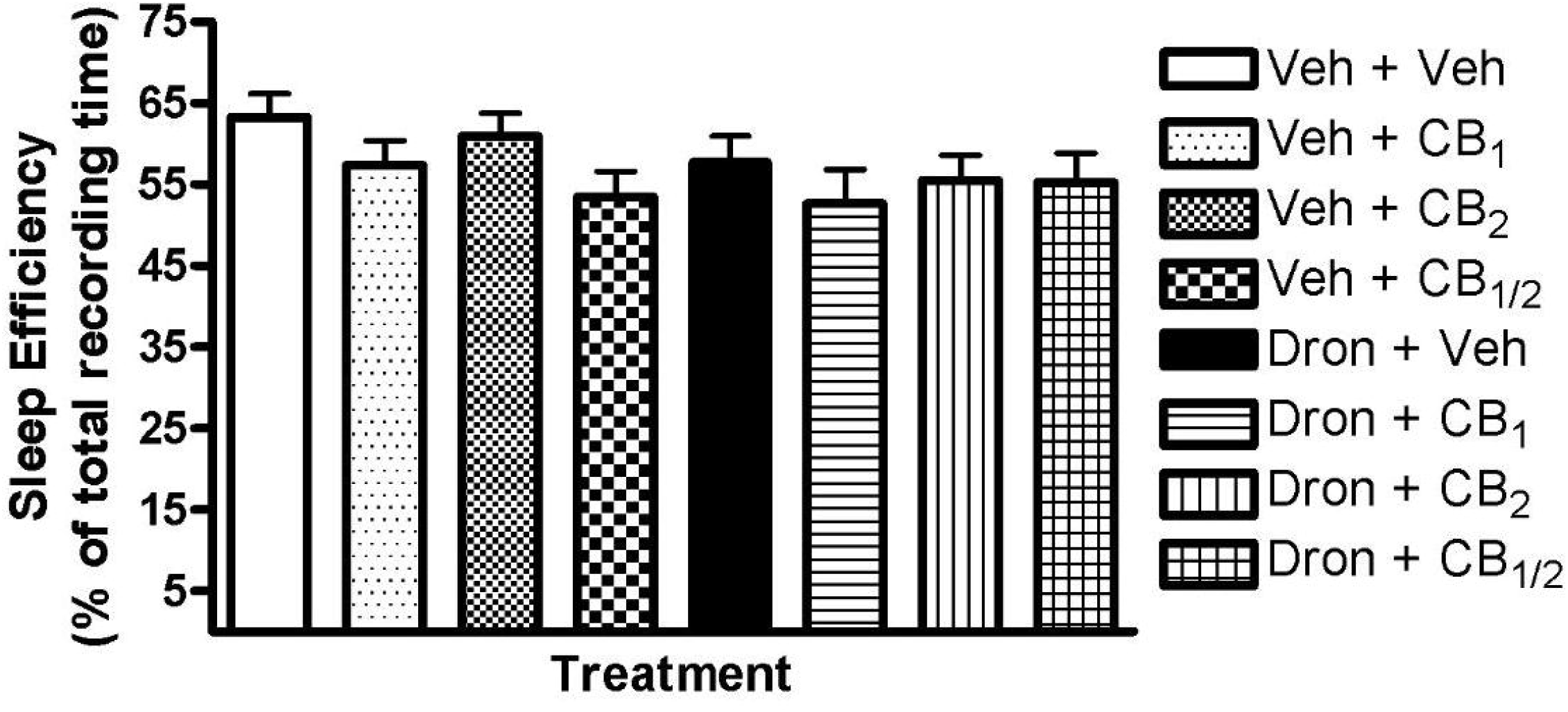
Sleep efficiency quantified as a percentage of time spent asleep from 6 hour recordings of conscious chronically-instrumented rat experiments. Vehicle (25% DMSO in PBS) or dronabinol (10 mg/kg) was injected IP in combination with vehicle or CB_1_ receptor (AM 251, 5 mg/kg) or CB_2_ receptor (AM 630, 5 mg/kg) antagonist, or both. There were no significant main effects. Data (mean ± SEM) were analyzed using mixed model analysis with repeated/fixed measures (CB agonist and CB antagonist).

**Figure 2.**
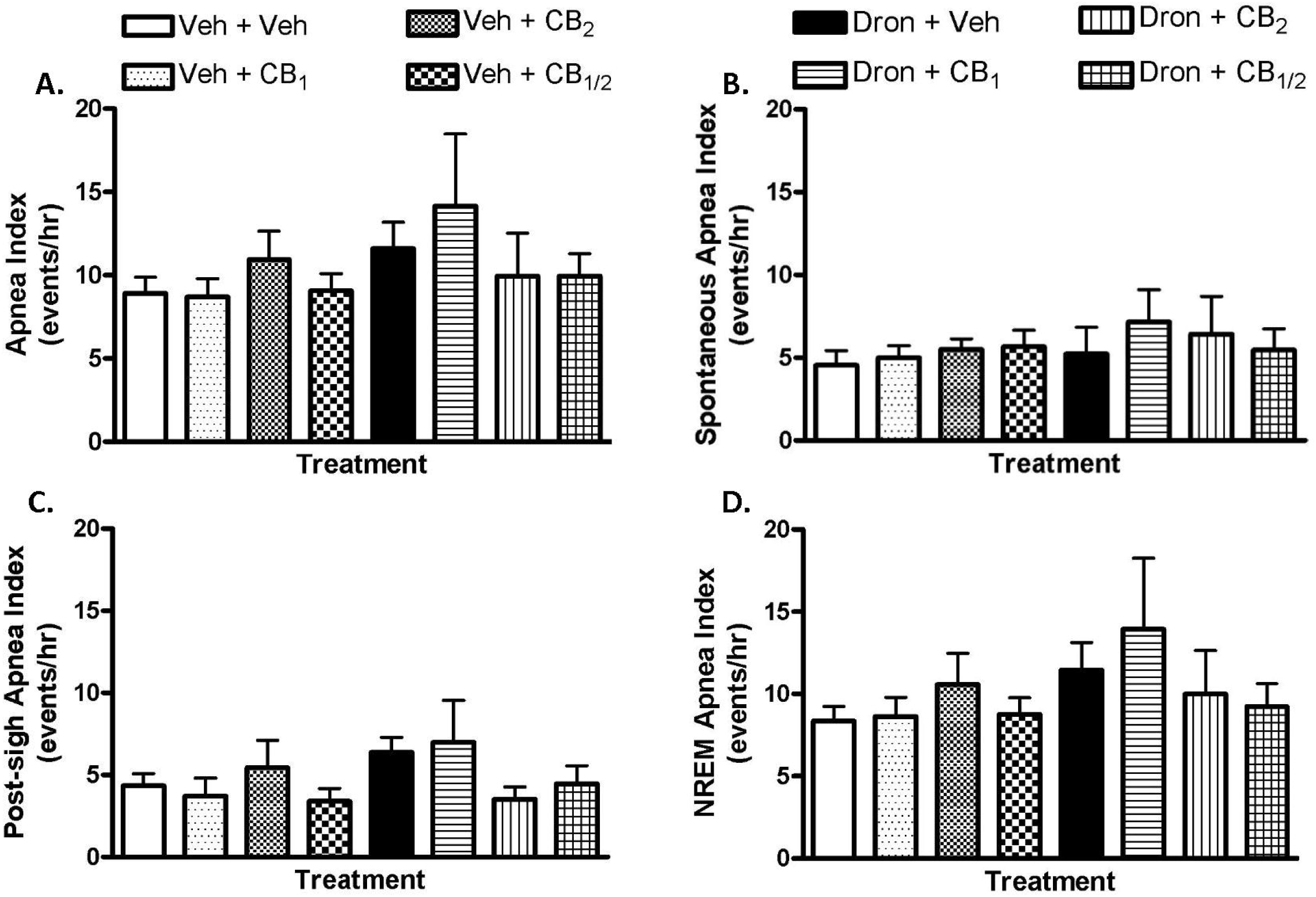
Apnea (**A**), spontaneous apnea (**B**), post-sigh apnea (**C**) and NREM apnea (**D**) indices quantified from 6 hour recordings of conscious chronically-instrumented rat experiments. Indices were quantified as events/hr. Vehicle (25% DMSO in PBS) or dronabinol (10 mg/kg) was injected IP in combination with vehicle (solid bars) or CB_1_ receptor (AM 251, 5 mg/kg) or CB_2_ receptor (AM 630, 5 mg/kg) antagonist, or both. There were no significant main effects. Data (mean ± SEM) were analyzed using mixed model analysis with repeated/fixed measures (CB agonist and CB antagonist).

**Figure 3.**
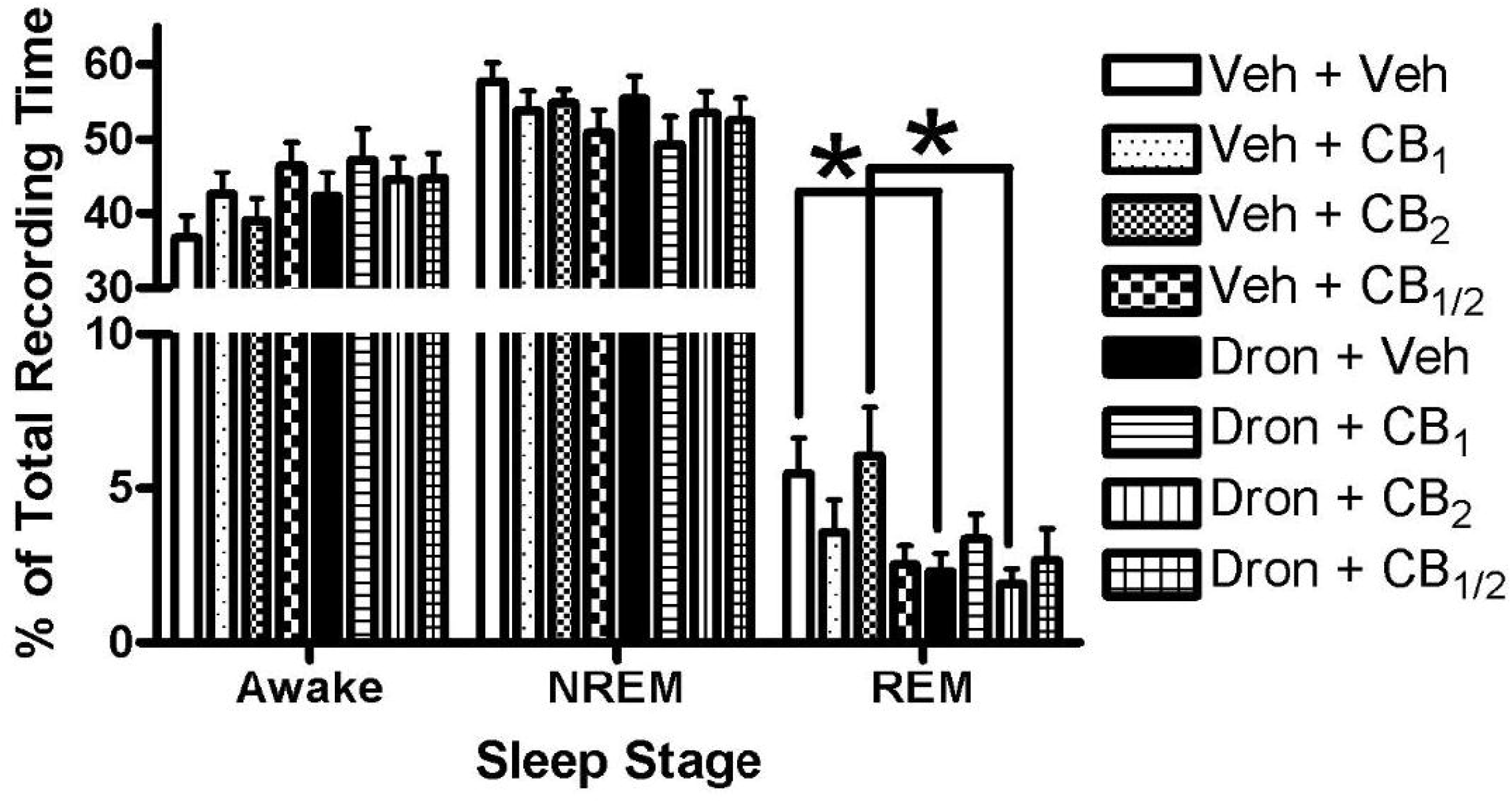
Awake time (**left**), and NREM (**center**) and REM (**right**) sleep as a percentage of total recording time quantified from 6 hour recordings of conscious chronically-instrumented rat experiments.. Vehicle (25% DMSO in PBS) or dronabinol (10 mg/kg) was injected IP in combination with vehicle or CB_1_ receptor (AM 251, 5 mg/kg) or CB_2_ receptor (AM 630, 5 mg/kg) antagonist, or both. Dronabinol and a combination of dronabinol and CB_2_ antagonist significantly reduced REM sleep. Data (mean ± SEM) were analyzed using mixed model analysis with repeated/fixed measures (CB agonist and CB antagonist) followed by post hoc multiple comparison tests with Sidak’s correction if there were significant main effects or a significant interaction of main effects. \**p* < 0.05.

The mixed model analysis revealed a significant effect of antagonist treatment (*F*_3, 64.41_ = 2.86, p = 0.04) on sleep efficiency (Fig. 1); however, post hoc analysis revealed no significant differences among the antagonist treatment groups (p > 0.05). There were no significant main effects (p > 0.05) on overall apnea index (Fig. 2A), spontaneous apnea index (Fig. 2B), post-sigh apnea index (Fig. 2C), and NREM apnea index (Fig. 2D). These results are in opposition with previous research showing an effect of dronabinol in 100% DMSO (1 ml) on sleep efficiency and apnea frequency [17,18].

No treatment effects were observed for time spent in NREM sleep (Fig. 3). Antagonist treatment had an effect (*F*_3, 65.43_ = 2.84, p < 0.05) on time spent awake, but post hoc analysis revealed no differences among the antagonist treatments (p > 0.05). There was significant agonist/antagonist interaction (*F*_3, 77.00_ = 3.68, p = 0.02) observed for REM sleep time. Post hoc analysis revealed that rats receiving dronabinol alone (2.31 ± 0.58%, N = 12) or dronabinol and CB_2_ antagonist (2.68 ± 1.02%, N = 12) had significantly (p< 0.01) decreased REM sleep compared to vehicle only (5.49 ± 1.13%, N = 12) or CB_2_ antagonist only (6.06 ± 1.56%, N = 12), respectively. These results are similar with previous research [17,18].

## Discussion

The major findings of the present study are: (a) dronabinol in 25% DMSO failed to suppress apneas or to decrease sleep efficiency (Figs. 1 & 2); and (B) dronabinol in 25% DMSO decreased REM sleep (Fig. 3), which is in contrast and in line, respectively, with previously reported results [17,18]. The only difference in experimental protocol between this study and previously reported studies was the concentration of DMSO used to dissolve dronabinol.

Dronabinol, a synthetic version of Δ9-THC, is a lipophilic substance that has been previously used to suppress apneas in preclinical studies [17,18] by a mechanism that involves modulation of vagus nerve activity via CB receptors on nodose ganglia [19,20]. In those preclinical studies, dronabinol was dissolved in undiluted DMSO since DMSO was known to increase bioavailability of the lipophilic drugs [3–5]. Although undiluted DMSO alone did not alter apnea expression in these studies [4, 8], DMSO may have altered the effects of dronabinol, since it is known to block fast axonal transport in the vagus nerve. Still, it is important to note that dronabinol dissolved in sesame oil rather than DMSO did reduce apnea frequency in two clinical trials in patients with obstructive sleep apnea syndrome [25,26].

In rats, dronabinol impacted sleep efficiency and apnea expression only when dissolved in 100% DMSO [17,18]. Decreasing the concentration of DMSO, as in the present study, eliminated the apnea suppressive effects. However, vehicle controls in those previous studies and in the present study had similar sleep efficiencies and apnea indices, arguing against the effect of DMSO alone on these parameters. The simplest explanation for this effect was that the increased concentration of DMSO increased the absorption and bioavailability of dronabinol [3]. Increased absorption and bioavailability of dronabinol could increase activation of CB receptors on the nodose ganglia, which play a part in apnea suppression [19,20]. However, we cannot rule out that DMSO potentiated the effects of dronabinol, by either modulating vagal nerve activity [9,27], or by modulating pulmonary ventilation [12], or both. Another plausible explanation is that dronabinol had increased access to the brain because DMSO decreased the integrity of the BBB [8]. The BBB is efficient at limiting the transport of Δ9-THC into the brain [28], thus decreased BBB integrity may increase the amount of dronabinol available to the brain and thus, modulation of breathing via centrally-located CB receptors [29]. However, a recent study that injected dronabinol into the brain demonstrated no effect on reflex apneas in anesthetized rats [21].

In contrast to the effects on apnea and sleep efficiency, the only measured effect of dronabinol in 25% DMSO was reduced REM sleep, which is in line with previous work using 100% DMSO [17,18]. Although the amount of REM sleep in vehicle treated rats was similar for 25% and 100% DMSO, the effects of dronabinol were larger in the 100% DMSO formulation compared to the 25% DMSO. This argues that 100% DMSO increased the bioavailability of dronabinol in comparison to 25% DMSO, since CB receptors located in the brain play a role in sleep regulation in rats [30], though both DMSO formulations allowed for enough dronabinol to be available to decrease REM sleep. Vagal nerve activity also has been implicated in sleep regulation [31,32], so the combination of dronabinol and 100% DMSO, with their known effects on the vagus nerve as previously discussed, might decrease REM sleep to a greater extent.

In conclusion, we show that dronabinol, a non-specific cannabinoid receptor agonist shown to suppress sleep-related apneas and REM sleep, does not suppress sleep-related apneas 25% DMSO vehicle. This adds to the growing literature that DMSO is not simply a compound used to dissolve polar and nonpolar compounds, but is a compound with its own innate biological activity.

## Acknowledgments

This was supported by the National Institutes of Health Grant 1UM1HL112856.

## Compliance with Ethical Standards

### Funding

This study was funded by National Institutes of Health (Grant 1UM1HL112856).

### Conflict of Interest

Michael W. Calik, PhD, has no conflicts of interest to disclose. David W. Carley, PhD, has conflicts of interest: stock/stockholder, royalties, and intellectual property rights from RespireRx (formerly Cortex Pharmaceuticals).

### Ethical approval

All applicable international, national, and/or institutional guidelines for the care and use of animals were followed.

